# Immunological responses to hydrogel-aided induced pluripotent stem cell-derived dopaminergic progenitor transplants in immunodeficient versus cyclosporine immunosuppressed rats

**DOI:** 10.64898/2026.06.09.731056

**Authors:** Giulia Comini, Tommy Patton, Nicola J Drummond, Michela Barbato, Oliver Treacy, Aideen E Ryan, Tilo Kunath, Eilís Dowd

## Abstract

The success of stem cell-derived brain repair for Parkinson’s is limited by the variable survival and poor maturation of dopaminergic progenitors after transplantation into the Parkinsonian brain. One approach that has been developed to improve this is engraftment of the cells within a neurotrophin-enriched collagen hydrogel. Although this has been shown to improve progenitor survival and maturation in athymic nude rats, the same beneficial effects of the hydrogel were not seen in cyclosporine immunosuppressed rats. To determine the reasons for these differences, the aim of this study was to assess the local and systemic immune responses to progenitor transplantation in these two recipient groups. To do so, human induced pluripotent stem cell-derived dopaminergic progenitors were transplanted into 6-hydroxydopamine-lesioned striatum of athymic or cyclosporine immunosuppressed rats. The cells were transplanted either alone, with the neurotrophins GDNF and BDNF, in an unloaded collagen hydrogel, or in a neurotrophin-loaded collagen hydrogel. *Post-mortem* assessment included both graft site and blood analysis of immune cell populations. As expected, nude rats showed a pronounced innate immune cell response at the graft site but no T-cell recruitment or activation locally or systemically. In contrast, while the immunosuppressed rats also showed the expected innate immune cells response to the transplant, there was also infiltration of CD4^+^ and CD8^+^ T cells at the site of transplantation as well as circulating activated T-cells. Thus, this study suggests that the benefits of the hydrogel that were seen in the athymic nude rats did not manifest in the cyclosporine immunosuppressed rats due to incomplete immunosupression. This study shows the importance of careful optimisation of the immunosuppressive regime chosen before xenotransplantation experiments.

## Introduction

Over the past two decades, numerous preclinical studies have established the safety and efficacy of human stem cell-derived dopaminergic grafts, leading to several ongoing clinical trials (Kyoto Trial: UMIN000033564; BlueRock: NCT04802733; STEM-PD: NCT05635409; S. Biomedics: NCT05887466; Aspen Neuro: ANPD001) and the approval of the first ever induced pluripotent stem cell (iPSC)-based therapy for Parkinson’s disease (PD) in Japan (Amchepry®; Sumitomo Pharma and Racthera). The success of these efforts depends critically on the transplanted progenitors’ ability to survive, mature, and functionally integrate within the Parkinsonian brain. However, preclinical studies suggest that these outcomes remain sub-optimal: graft survival is highly variable, ranging from less than 1% to 500% of transplanted cells (median: 51%), and the dopaminergic differentiation efficiency of transplanted dopaminergic progenitors (DAPs) is poor, varying from 0% to 46% (median: 3% across models and only 1% in non-human primates) (Comini & Dowd, 2025). To fully realize the therapeutic potential of this approach, there is a critical need to improve transplantation outcomes, particularly cell survival and dopaminergic maturation efficiency *in situ*.

Poor survival and maturation may result from multiple factors, including growth factor deprivation in the adult Parkinsonian brain, hostile immune responses, and the absence of a structural matrix for cell adhesion post-transplantation (Abeliovich et al., 2007; Duan et al., 1995). In this regard, biomaterials have been used as a physical scaffold to which cells can adhere thereby mimicking the extracellular matrix which is lost after dissociation of stem cell-derived dopaminergic progenitors (DAPs) from coated cell culture plates (Béduer et al., 2015, Newland et al., 2015b). To address the issue of neurotrophin-deprivation that the cells experience once transplanted, biomaterials have been functionalised with neurotrophic growth factors creating a favourable microenvironment for cell survival and differentiation (Adil et al., 2017, Hoban et al., 2013, Cabré et al., 2021a, Meng et al., 2020). Finally, certain biomaterials can act as a physical barrier and protect the transplanted cells from the host neuroimmune response (Hoban et al., 2013, Moriarty et al., 2017).

We recently demonstrated that a neurotrophin-enriched collagen hydrogel can enhance survival (8-fold) and maturation (16-fold) of human iPSC-DAPs after intra-striatal transplantation in athymic nude rats compared to when iPSC-DAPs were transplanted alone (Comini et al., 2024). However, this beneficial effect was not observed when the same experiment was conducted in cyclosporine A (CsA) immunosuppressed rats. Athymic nude rats and CsA immunosuppressed rats differ fundamentally in their immune systems. Nude rats lack the thymus due to a FOXN1 mutation, resulting in deficient and immature T cells (Schuddekopf et al., 1996, Hirasawa et al., 1998, Hanes, 2006). In contrast, cyclosporine functions as a T-cell inhibitor by inhibiting calcineurin/NFATc-mediated transcription of IL-2 which is required for T-cell activation (Lee et al., 2023).

Considering these factors, the study presented in the current manuscript examined the immunological responses to iPSC-DAP transplantation in both athymic nude and cyclosporine immunosuppressed rats to determine the differences between these recipient groups.

## Methods

### Ethical statements

All procedures involving animals were approved by the Animal Care and Research Ethics Committee at the University of Galway; were completed under Project and Individual Authorisations issued by the Irish Government via the Health Products Regulatory Authority; and were carried out in compliance with the European Union Directive 2010/63/EU and the Irish Statutory Instrument No. 543 of 2012. Immunosuppressed and nude rats were housed in pairs in individually ventilated cages (IVCs). In each cage, standard bedding material (3Rs lab basic bedding) and sizzle-nest and circular hollow plastic tunnels were provided as environmental enrichment. Animals were kept on a 12:12 h light:dark cycle (lights on at 07:00 am), at 19-23°C, with relative humidity levels maintained between 40 and 70%. For the duration of the experiments, rats were allowed food and water *ad libitum*. All procedures involving human cells were approved by the relevant Human Ethics Committees at the University of Edinburgh and the University of Galway.

### Experimental design

36 female athymic nude rats (Hsd:RH-*Foxn1*^rnu^; Envigo UK) and 36 female SD rats (Charles River, UK) received unilateral injection of 6-hydroxydopamine (6-OHDA) at medial forebrain bundle (MFB) coordinates. Two weeks later, the rats were randomly divided into 4 groups (Cells Alone; Cells & NTFs; Cells & Hydrogel; Cells & NTFs & Hydrogel) and given the corresponding intra-striatal transplant. CsA treatment for SD rats consisted of subcutaneous injections of 10 mg/kg of CsA starting from 24h before transplantation. In all groups, rats received 300,000 human iPSC-DAPs (from the NAS2 cell line at Day 16 of a midbrain floorplate differentiation protocol (Nolbrant et al., 2017; Drummond et al., 2020). Rats were then sacrificed and brains and whole blood were collected at day 1, 4 and 7 (n=3 per group, per time-point) after transplantation for *post-mortem* evaluation of adaptive and innate immune system reaction to the transplants through immunostaining and flow cytometry analysis (Fig. 1).

**Figure 1.**
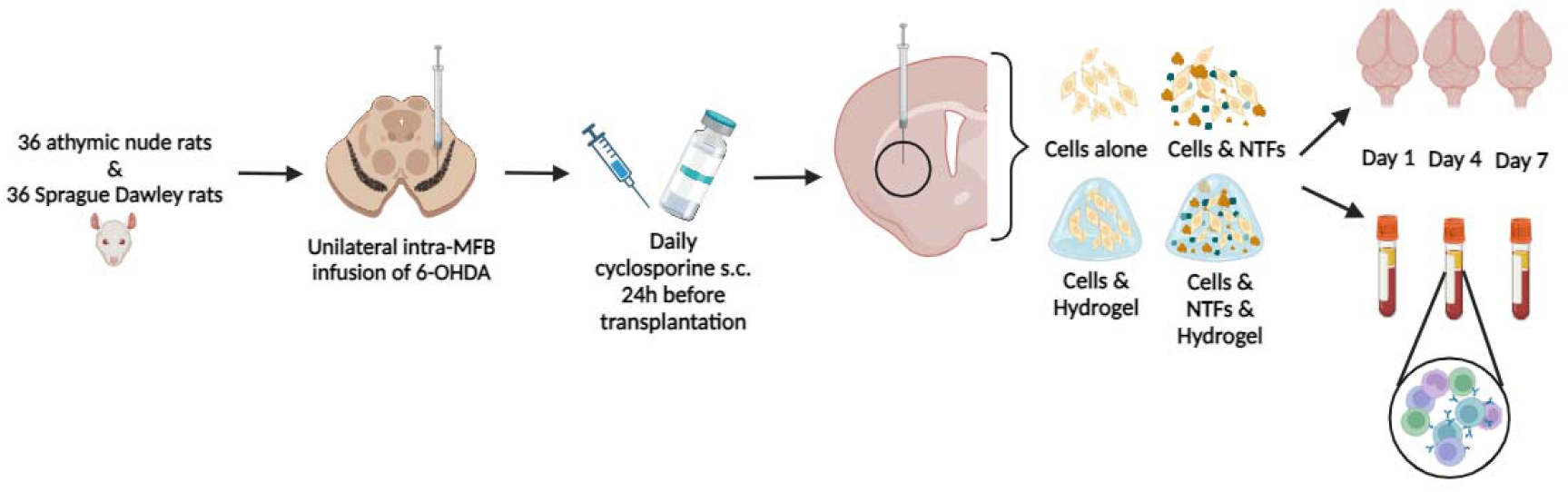
Schematic representation of the experimental design. iPSC-DAPs were transplanted either alone (300,000 cells/6 µl), with NTFs (GDNF 500 ng and BDNF 1000 ng), encapsulated in a collagen hydrogel or in a NTF-enriched collagen hydrogel in the brain of either 6-OHDA lesioned female athymic nude rats or 6-OHDA lesioned female SD rats immunosuppressed with daily subcutaneous CsA injections (10 mg/kg) from 24 hours prior to transplantation. *Post-mortem* assessments were carried out at 1, 4 and 7 days post-grafting and consisted of immunohistochemical analysis along with flow cytometry evaluation of the T-cell populations.

### Culture and differentiation of iPSC-DAPs

Human NAS2 iPSCs (Devine et al., 2011) were maintained on laminin-521-coated plates in iPSC brew medium. For neuronal patterning (day 0), cells were dissociated using 0.5 mM EDTA, seeded onto laminin-111-coated wells, and cultured in neural induction medium (NIM) supplemented with Y-27632, SB431542, LDN-193189, SHH-C24II, and CHIR99021. Medium was replaced on day 2. From days 4-9, cells were maintained in neural proliferation medium (NPM) containing SB431542, LDN-193189, SHH-C24II, and CHIR99021, with medium changes every 2-3 days. On day 9, the medium was switched to NPM supplemented with FGF8b and heparin solution. On day 11, cells were dissociated with accutase and re-plated onto laminin-111-coated wells in neural differentiation medium (NDM) supplemented with Y-27632, BDNF, GDNF, FGF8b, ascorbic acid (AA), and heparin solution to promote dopaminergic specification. Medium was changed on day 14. By day 16, cells were prepared for transplantation. Detailed protocol: dx.doi.org/10.17504/protocols.io.bddpi25

### Cell suspension preparation

On day 16 of differentiation, cells were dissociated from wells using accutase and centrifuged at 1100 rpm for 5 minutes. The cell pellet was resuspended in 1.2 ml of NDM, and a 10 µl aliquot was taken for cell counting. Following quantification, cells were centrifuged again and resuspended in NDM at a final density of 300,000 cells/6 µl for hydrogel preparation and transplantation.

### Hydrogel fabrication

To prevent early gelation during collagen hydrogel preparation, all materials were maintained on ice throughout the procedure. Initially, type 1 atelocollagen (5 mg/ml) was neutralized to pH 7 using 1M NaOH. The NTF-enriched collagen hydrogels for transplantation were then prepared by thoroughly combining neutralized collagen (40%), cell suspension in neural differentiation media (NDM) (30%), 4s-StarPEG crosslinker at 4 mg/mL in PBS (10%), and neurotrophic factors consisting of GDNF (500 ng) and BDNF (1000 ng) in PBS (20%). In control groups lacking certain components, NDM was used as a substitute to maintain equivalent volumes.

### Surgeries

All surgical procedures were conducted using isoflurane anaesthesia (5% in O□ for induction, 2% in O□ for maintenance) with animals positioned in a stereotaxic frame. The nose bar was set at −4.5 for intra-MFB procedures or −2.3 for intra-striatal procedures. During lesioning, 3 µl of 6-OHDA solution (12 mg in 3 ml) was unilaterally delivered into the MFB at stereotaxic coordinates: Anterior-Posterior (AP) −4.0, Medial-Lateral (ML) −1.3 relative to bregma, and Dorsal-Ventral (DV) −7.0 below the dura in SD rats and AP −3.9, ML −1.2 and DV −7.3 in athymic nude rats. For transplantation procedures, 6 µl of cell suspension (with or without collagen hydrogel and NTFs) was infused into the striatum at coordinates: AP 0.0, ML −3.7 from bregma, DV −5.0 below the dura in SD rats and AP +1.3, ML −2.6, DV −5 in athymic nude rats, and the cannula level was raised slightly every 2 minutes. All infusions were administered at 1 µl/min, followed by an additional 2-minute waiting period to facilitate diffusion.

### Immunohistochemistry (IHC)

Animals were euthanized using terminal anaesthesia (pentobarbital 140 mg/kg intraperitoneal) followed by transcardial perfusion-fixation with heparinized saline and 4% paraformaldehyde (PFA). Extracted brains were fixed in 4% PFA overnight, then transferred to 25% sucrose-azide solution the following day for long-term preservation. Coronal brain sections (30 µm) were prepared using a freezing sledge microtome, and free-floating immunohistochemistry was performed to detect human nuclei (HuNu, not shown), T-helper cells (CD4), and cytotoxic T cells (CD8), activated microglia (OX-42), reactive astrocytes (GFAP).

### Peripheral blood mononuclear cells (PBMCs) isolation

PBMCs were isolated from whole blood obtained during transcardial perfusion prior to fixation. Blood was collected in K2EDTA-coated BD Vacutainer^®^ tubes to prevent coagulation and processed immediately for PBMCs extraction. The isolation protocol involved layering blood onto 4 ml of Ficoll (Sigma) in a centrifuge tube, followed by centrifugation at 2,000 rpm for 20 minutes to separate blood components by density. After discarding the upper plasma layer, PBMCs were collected from the interphase layer (buffy coat) and transferred to a tube containing sterile PBS. The cells were washed three times by centrifugation at 1,200 rpm for 5 minutes, with resuspension in fresh PBS after each spin. PBMC yield was determined by cell counting: 10 µl of cell suspension was mixed with an equal volume of Trypan Blue^®^, and 10 µl of this mixture was applied to a hemocytometer for manual counting under microscopy. PBMCs were subsequently stained to assess the presence of T-helper cells (PE/Cyanine 7 anti-rat CD4), CD8-positive cells (PerCP anti-rat CD8a), activated T-cells (PE anti-rat CD25) and NK cells (FITC anti rat CD335).

### Flow cytometry

Flow cytometry was conducted to evaluate T-cell and NK cell populations in blood samples. Briefly, 100,000 PBMCs were placed into individual V-bottom wells of a 96-well plate. Following centrifugation for 5 minutes, the supernatant was aspirated, leaving cells pelleted at the well bottom. For immunostaining, appropriate antibodies (listed in Table 1) were diluted in 50 µl of FACS buffer (DPBS with 2% FBS and 0.1% NaN□), added to the wells, and incubated for 15 minutes at 4°C while protected from light. Cells were washed twice by centrifugation with 100 µl of FACS buffer added after each spin. Finally, stained cells were resuspended in 200 µl of FACS buffer and transferred to 5 ml tubes, which were maintained on ice and shielded from direct light until analysis. Samples were analysed using a Cytek® Northern Lights Flow Cytometer and data was analysed using FlowJo® analysis software version 10 (Tree Star, Inc., Ashland, OR, USA).

**Table 1.** Antibodies and relative experimental concentrations used for flow cytometry.

| **Antibody** | **Target** | **Clone** | **Source** | **Concentration** |
| --- | --- | --- | --- | --- |
| ***FITC anti-rat CD335*** | NK p46 | WEN23 | Biolegend;<br>250806 | 10 µg/ml |
| ***PE/Cyanine 7 anti-rat CD4*** | T helper cells | W3/25 | Biolegend;<br>201516 | 1.28 µg/ml |
| ***PerCP anti-rat CD8α*** | CD8-positive cells | OX-8 | Biolegend;<br>201712 | 6 µg/ml |
| ***PE anti-rat CD25*** | Activated T cells | OX-39 | Biologend;<br>202105 | 4 µg/ml |

### Data and statistical analyses

All analyses were carried out by researchers blinded to the treatment of the animals. Immunostained sections were captured by an Olympus VS120 Virtual Slide Microscope and analysed using ImageJ software (U.S. National Institutes of Health). For qualitative observation, the time points were kept separate, while for statistical analysis of flow cytometry results and volumetric quantification, the three time point were collapsed together. All graphs show individual data points with mean ± standard error of the mean (SEM). Statistical analysis was performed utilizing GraphPad software (La Jolla, CA, USA) and using 1-way analysis of variance (ANOVA) with Tukey’s *post hoc*. Any other tests used are mentioned in the text.

## Results

### Adaptive immune system response to the transplant

Firstly, qualitative observation of the human nuclear staining in athymic nude rats (Fig. 2A) and in CsA suppressed rats (Fig. 2B) was performed to assess the presence of surviving transplants before studying the immunological profiles in the two strains. Grafts were detected across the four experimental groups and at every time point.

**Figure 2.**
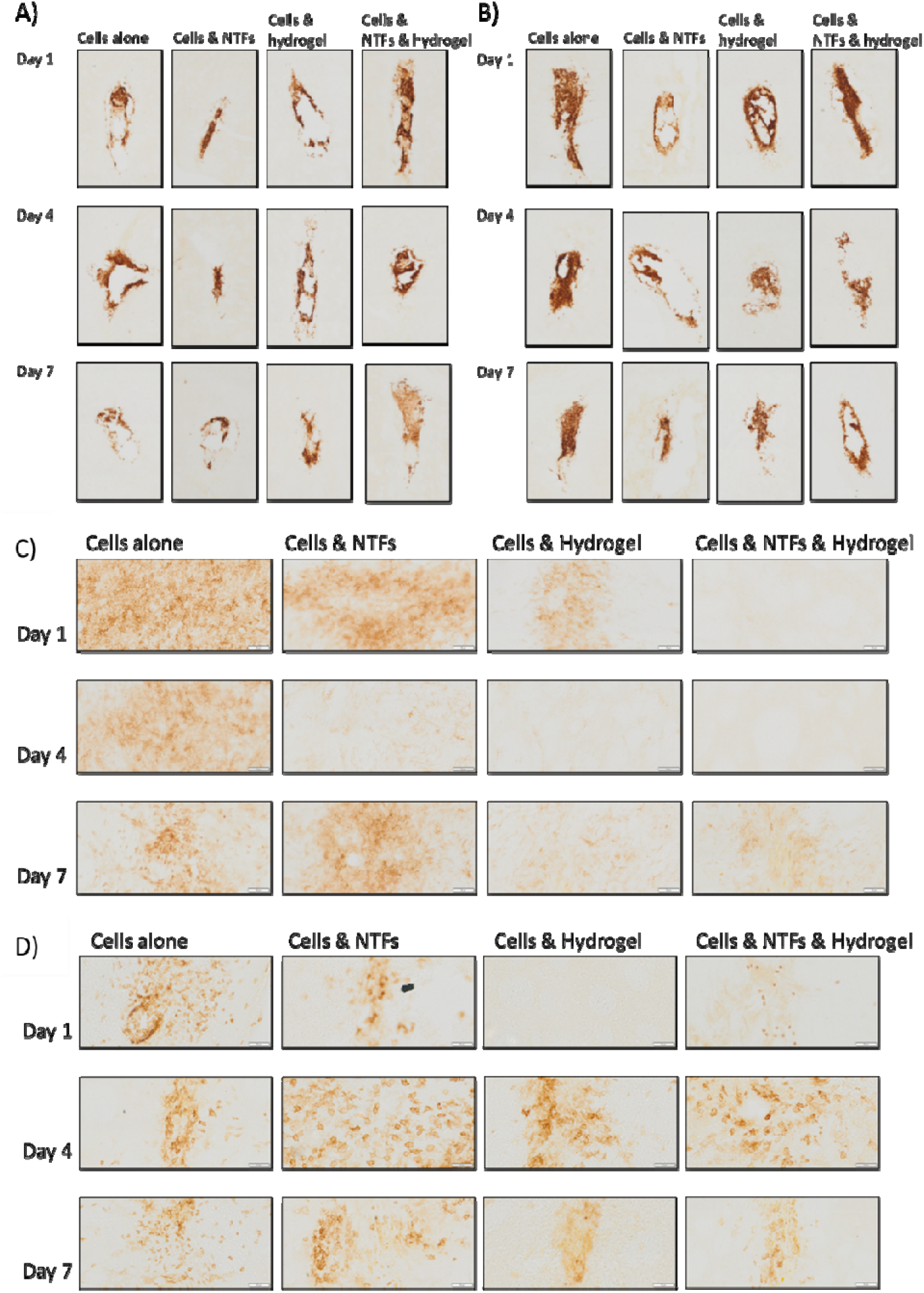
Human nuclear staining and adaptive immune system response to the transplant in CsA treated rats. Human nuclear staining of the striatal sections confirmed the presence of surviving iPSC-derived grafts both in nude rats (A) and CsA immunosuppressed SD rats (B) in all treatment groups and at all time points. Immunostaining for CD4^+^ T-helper (C) and CD8^+^ (D) cells on striatal sections of CsA immunosuppressed SD rats revealed infiltration of T-cells at the site of transplantation across all groups and all time points.

Adaptive immune system activation following iPSC-DAPs transplantation was evaluated by immunostaining for T-helper (CD4^+^) and CD8-positive (CD8^+^) cells in striatal sections at the graft site. CD4^+^ immunoreactivity was absent in athymic nude rats, as expected (not shown). However, CD4^+^ cells were detected in immunosuppressed rats across all time points and experimental conditions (Fig. 2C). CD4^+^ staining was evident in 3/9 rats receiving cells alone, 4/9 with NTFs, 2/9 with unloaded collagen hydrogel encapsulation, and 1/9 with NTF-enriched collagen hydrogel.

Similarly, CD8^+^ immunoreactivity was absent in athymic nude rats (not shown), but present in immunosuppressed animals across all groups and time points (Fig. 2D). CD8^+^ staining was observed in 4/9 rats with cells alone, 3/9 with NTFs, 2/9 with unloaded collagen hydrogel, and 2/9 with NTF-enriched collagen hydrogel.

### Innate immune system response to the transplant

The innate immune response following transplantation of iPSC-DAPs, either with or without NTFs and/or collagen hydrogel, was evaluated by assessing microgliosis (OX-42 marker) and astrocytosis (GFAP marker) immunostaining in striatal sections at the graft site.

In terms of microgliosis, OX-42 positive staining was observed at all post-transplantation time points in the striata of athymic nude rats (Fig. 4A), and a similar microgliotic response was detected in all experimental groups in CsA-treated rats (Fig. 4B). Similarly, mild astrocytic reactivity (GFAP-positive staining) was observed across all experimental groups in both athymic nude rats (Fig. 4C) and CsA-immunosuppressed rats, regardless of NTF or hydrogel inclusion (Fig. 4D).

### Flow cytometry analysis of circulating T-cells and NK cells

In parallel, flow cytometry analysis was performed to assess circulating T-cell populations in athymic nude rats and CsA immunosuppressed rats following iPSC-DAP transplantation. For each T-cell subset, total circulating CD4^+^ & CD8^+^ cells and activated CD4^+^ and CD8^+^ (CD4^+^CD25^+^ & CD8^+^CD25^+^) cells were quantified. CD4^+^ T-helper cells were present in the peripheral blood of athymic nude rats but were not activated (CD25^−^) (Fig. 3A&Ai). In contrast, CsA treated rats exhibited both circulating and activated (CD4^+^CD25^+^) T-helper cells phenotypes, although without significant intergroup differences (Fig. 3B&Bi).

**Figure 3.**
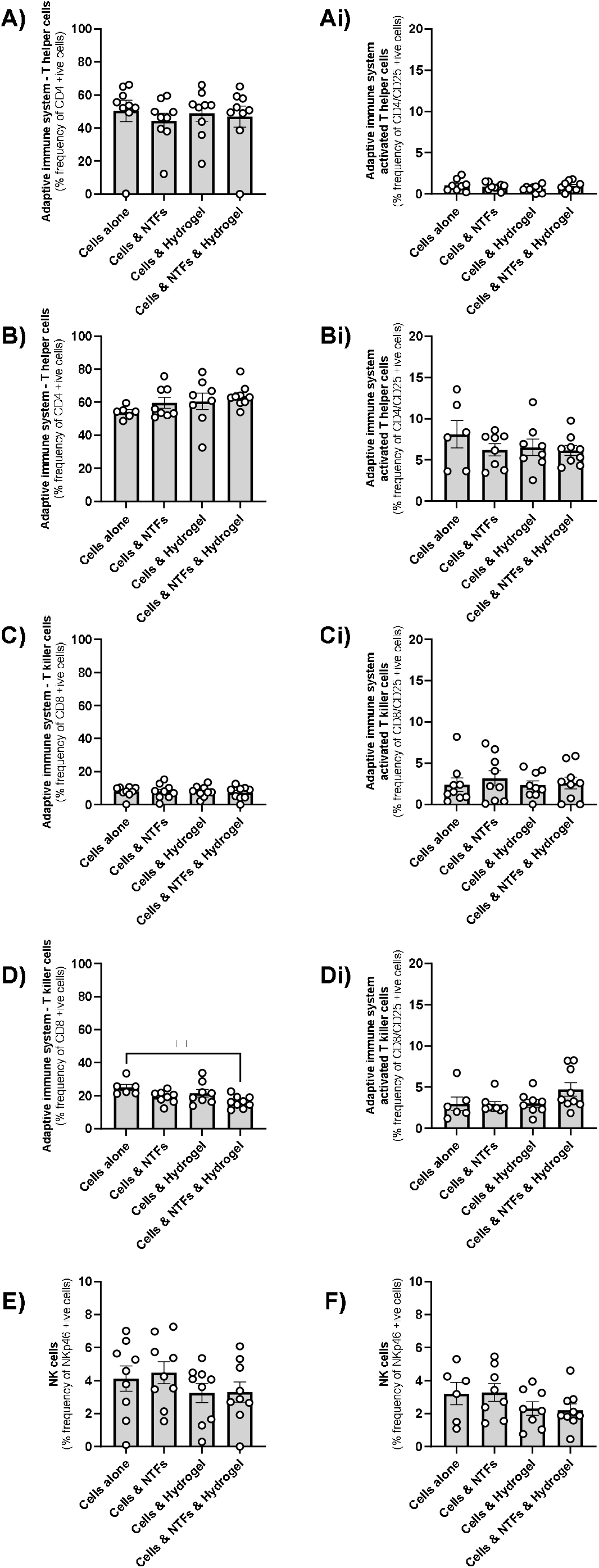
Study of the systemic immune system response to the transplant. Flow cytometry analysis for circulating (CD4^+^, CD8^+^) and activated (CD4^+^CD25^+^, CD8^+^CD25^+^) T-helper and CD8-positive cells; and NK cells (CD335^+^) was performed on the PBMCs isolated from the blood of athymic nude rats and CsA immunosuppressed rats in which iPSC-DAPs were transplanted either alone, with NTFs, in a hydrogel or in a NTF-enriched hydrogel. A) Circulating CD4^+^ cells in athymic nude rats, Ai) activated CD4^+^CD25^+^ cells in athymic nude rats, B) circulating CD4^+^ cells in CsA suppressed rats, Bi) activated CD4^+^CD25^+^ cells in CsA suppressed rats, C) circulating CD8^+^ cells in athymic nude rats, Ci) activated CD8^+^CD25^+^ cells in athymic nude rats, D) circulating CD8^+^ cells in CsA suppressed rats, Di) activated CD8^+^CD25^+^ cells in CsA suppressed rats, E) circulating CD335^+^ cells in athymic nude rats and F) circulating CD335^+^ cells in CsA suppressed rats. Data represent individual data points, are expressed as mean ± SEM and were analysed by one-way ANOVA and Tukey’s *post-hoc*.

**Figure 4.**
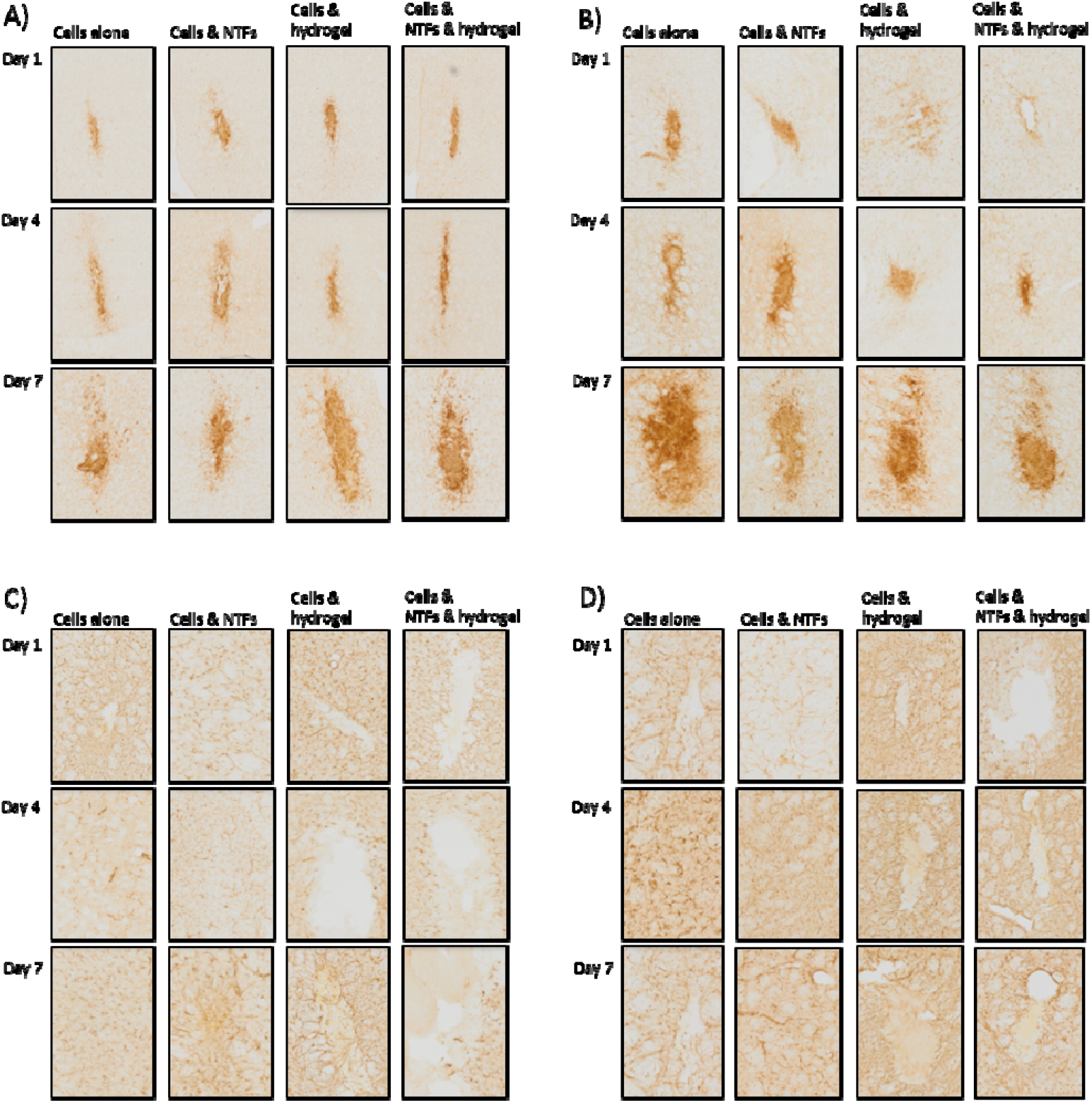
Study of the innate immune system response to the transplant. Immunostaining for microglia (OX-42 marker; A & B) and astrocytes (GFAP marker; C & D) was performed on the striatal sections of athymic nude rats (A & C) and CsA immunosuppressed rats (B & D) in which iPSC-DAPs were transplanted either alone, with NTFs, in a hydrogel or in a NTF-enriched hydrogel.

CD8^+^ cells were detected at low levels in athymic nude rats, with minimal activation (Fig. 3C&Ci). CsA treated rats showed higher frequencies of circulating CD8^+^ cells also with low activation levels. Notably, NTF-enriched hydrogel encapsulation significantly reduced circulating CD8^+^ cell frequency compared to cells transplanted alone (F_(3,27)_=4.37, **P<0.01) (Fig. 3D&Di).

Additionally, flow cytometry analysis was conducted to assess circulating NK cells (CD335□) in both rat strains. NK cells were detected in the blood of both athymic nude rats (Fig. 3E) and CsA suppressed rats (Fig. 3F). However, no significant differences were found among experimental groups in either strain.

## Discussion

Numerous clinical trials involving putaminal grafting of human ESC- or iPSC-DAPs to assess the safety of this approach in PD are currently ongoing. Moreover, the first iPSC-based therapy for patients has recently been approved in Japan. However, the success of these interventions depends critically on the efficient survival and maturation of transplanted cells in the brain. In this regard, a systematic review of the preclinical literature revealed considerable variability, with generally low survival and particularly limited differentiation of the progenitors following transplantation (median: 3%) (Comini & Dowd 2025).

In an effort to improve these outcomes, the potential of a NTFs-enriched collagen hydrogel to enhance the survival and maturation of iPSC-DAPs following transplantation into the brains of either athymic nude rats (T-cell deficient) or CsA suppressed rats (T-cell suppressed) was previously investigated. Interestingly, the hydrogel, either alone or loaded with NTFs, was able to increase the survival and maturation of transplanted cells in athymic nude rats (Comini et al., 2024). However, when the same experiment was performed in CsA suppressed rats, although no graft rejection was observed, the hydrogel did not confer the same beneficial effect (study not published).

Therefore, the main aim of the study presented in this manuscript was to profile the immunological status of athymic nude rats and CsA suppressed rats following transplantation of human iPSC-DAPs, administered either alone or in combination with NTFs and/or a collagen hydrogel, in order to identify possible reasons underlying the previously reported differences between the two strains.

Briefly, the adaptive immune response in the striatum was assessed, as well as in the peripheral blood of the host animals following transplantation. Given that athymic nude rats are a specific strain that lacks appropriate T-cell lymphocyte development due to the absence of a thymus (Brooks et al., 1980; Hanes, 2006), as expected, they showed low levels of circulating and activated T cells after transplantation. In parallel, fully immunocompetent SD rats were administered CsA, an immunosuppressive drug widely regarded as the gold standard in rodent studies, which acts by suppressing T-cell mediated responses. It is typically administered 24 hours prior to transplantation at a dose of 10 mg/kg and continued for the duration of the experiment to prevent graft rejection (Kriks et al., 2011; Kirkeby et al., 2012; Kikuchi et al., 2017b; Shrigley et al., 2021b).

T-cell specific immunostaining revealed infiltrating T-helper and cytotoxic T cells at the transplantation site in the brains of CsA immunosuppressed SD rats. To further investigate these findings, FACS analysis on PBMCs isolated from the blood of both strains at the time of sacrifice was performed. This showed elevated levels of activated T-helper cells and circulating cytotoxic T cells in CsA treated rats, whereas no such populations were detected in immunodeficient nude rats.

These findings indicate that SD rats were not fully immunosuppressed at the time of transplantation and that subcutaneous CsA administration at 10 mg/kg, initiated 24 hours prior to surgery, is not completely effective in suppressing T-cell mediated responses. Moreover, although collagen hydrogels have been proposed to act as a physical barrier between the host neuroimmune system and transplanted progenitors (Hoban et al., 2013; Cabré et al., 2021a), this protective effect may not extend to T cells, which have been shown to migrate through and infiltrate collagen matrices (Sadjadi et al., 2020). These findings may explain why the collagen hydrogel exerted beneficial effects in athymic nude rats but not in CsA treated rats, as previously observed.

In addition to the differences observed between the two strains, the use of nude animals as preclinical models as been shown to significantly improve transplantation outcomes, particularly in terms of *in situ* differentiation of stem cell-derived dopaminergic progenitors (Comini & Dowd 2025). This supports the hypothesis that T-cell deficient and T-cell suppressed models represent fundamentally different experimental conditions.

Furthermore, it is important to consider whether the use of T-cell deficient animals is as relevant for clinical translation as the use of immunosuppressed hosts. Transplantation in immunosuppressed animals is generally thought to more closely resemble allogeneic transplantation (donor-to-patient) in clinical settings, rather than autologous transplantation (patient-to-self).

Although both approaches offer distinct advantages, autologous transplantation presents significant logistical challenges, particularly in terms of scalability, as cells must be generated and quality-controlled individually for each patient. Conversely, allogeneic approaches allow for large-scale production, quality control, and storage of cell batches, making them more cost-effective, reliable, and standardised.

In light of these considerations, future studies should focus on evaluating strategies to improve the *in vivo* survival and differentiation of transplanted iPSC-DAPs for stem cell-based brain repair in immunosuppressed rather than immunodeficient models. Furthermore, given our finding that rats were not fully immunosuppressed at the time of transplantation following subcutaneous CsA administration (10 mg/kg, 24 hours prior to transplantation), the immunosuppressive regimen may require optimisation prior to xenotransplantation experiments.

## Supporting information

Supplementary, Gating Strategy

## Acknowledgements

This study was supported by research grants from the Michael J Fox Foundation for Parkinson’s Research (Grant Numbers: 17244 and 023410) and Science Foundation Ireland (Grant Number: 19/FFP/6554; 13/RC/2073_P2).

## Competing Interests

The authors declare no competing financial or non-financial interests.

## Data Availability

The complete datasets generated during the current study are available from the corresponding author on reasonable request.

## Author Contributions

ED led the design and implementation of the project and wrote the manuscript with GC; GC completed all aspects of the study helped by TP; AR and OT provided assistance for flow cytometry analysis. All authors have read and agreed to the published version of the manuscript.

## Notes

**Funding:** This study was supported by research grants from the Michael J Fox Foundation for Parkinson’s Research (Grant Numbers: 17244 and 023410) and Science Foundation Ireland (19/FFP/6554 and 13/RC/2073_P2).

### Competing Interest Statement

The authors have declared no competing interest.

