## Supplementary, Gating Strategy for "Immunological responses to hydrogel-aided induced pluripotent stem cell-derived dopaminergic progenitor transplants in immunodeficient versus cyclosporine immunosuppressed rats"

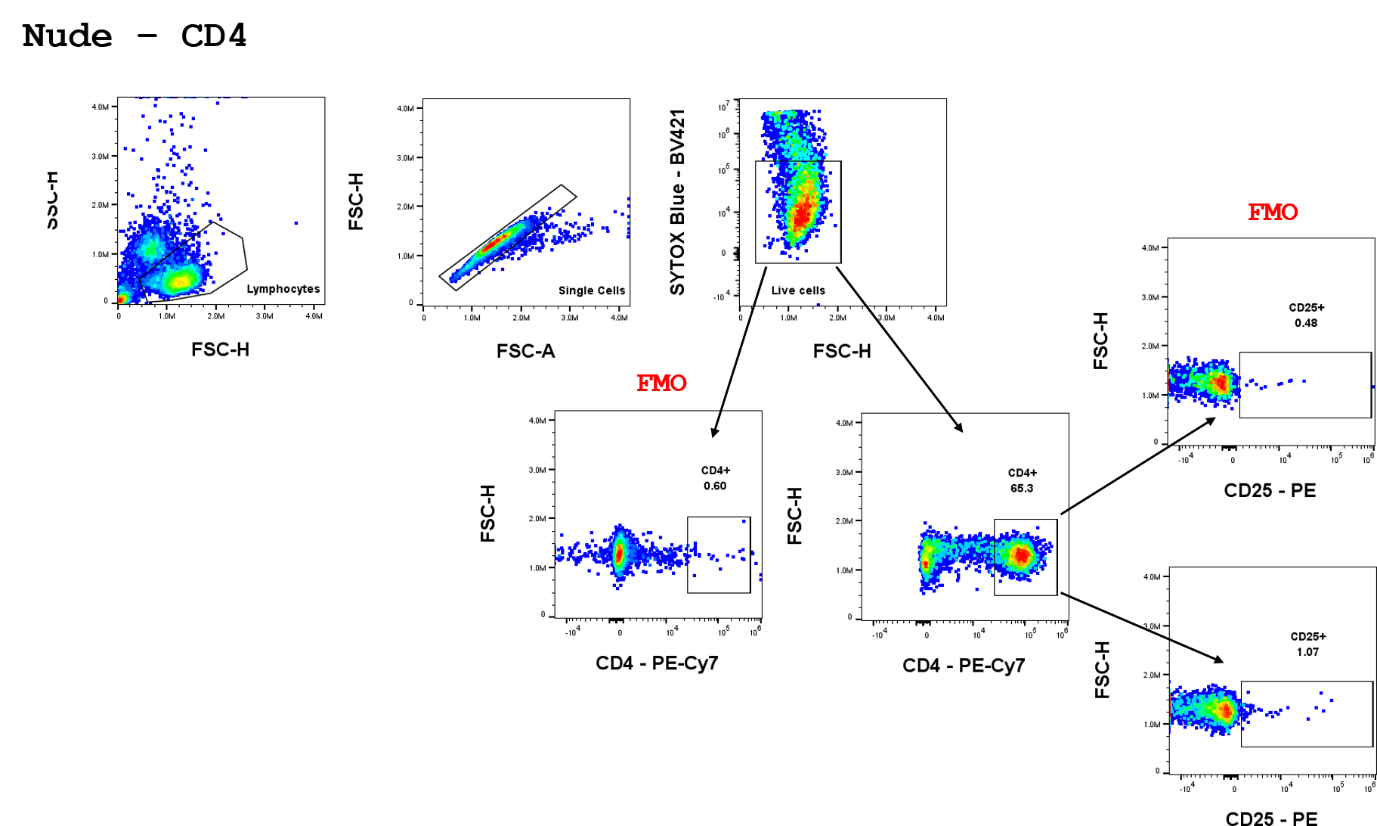

*Figure 1. Gating strategy for flow cytometric detection of CD4⁺ cells in PBMCs from nude rats.*

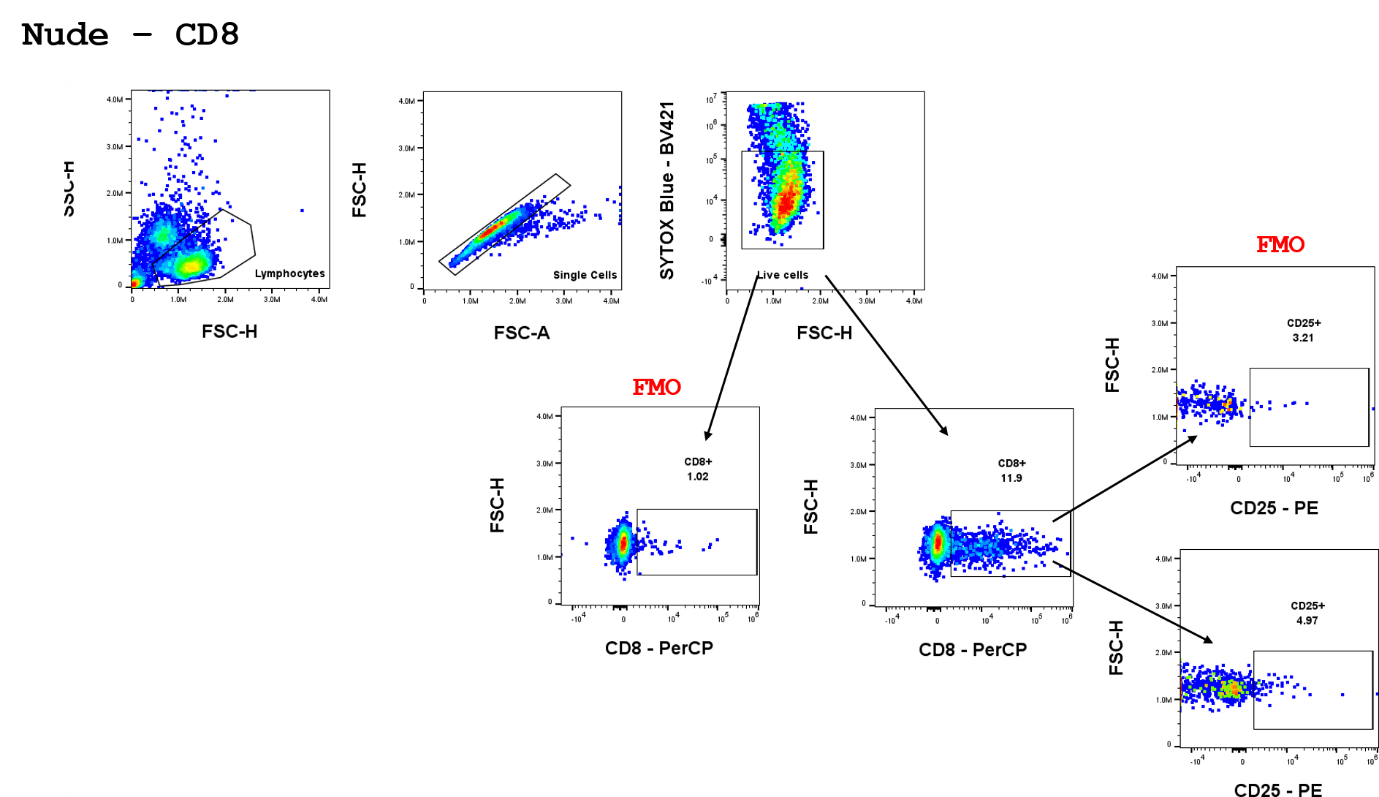

*Figure 2. Gating strategy for flow cytometric detection of CD8⁺ cells in PBMCs from nude rats.*

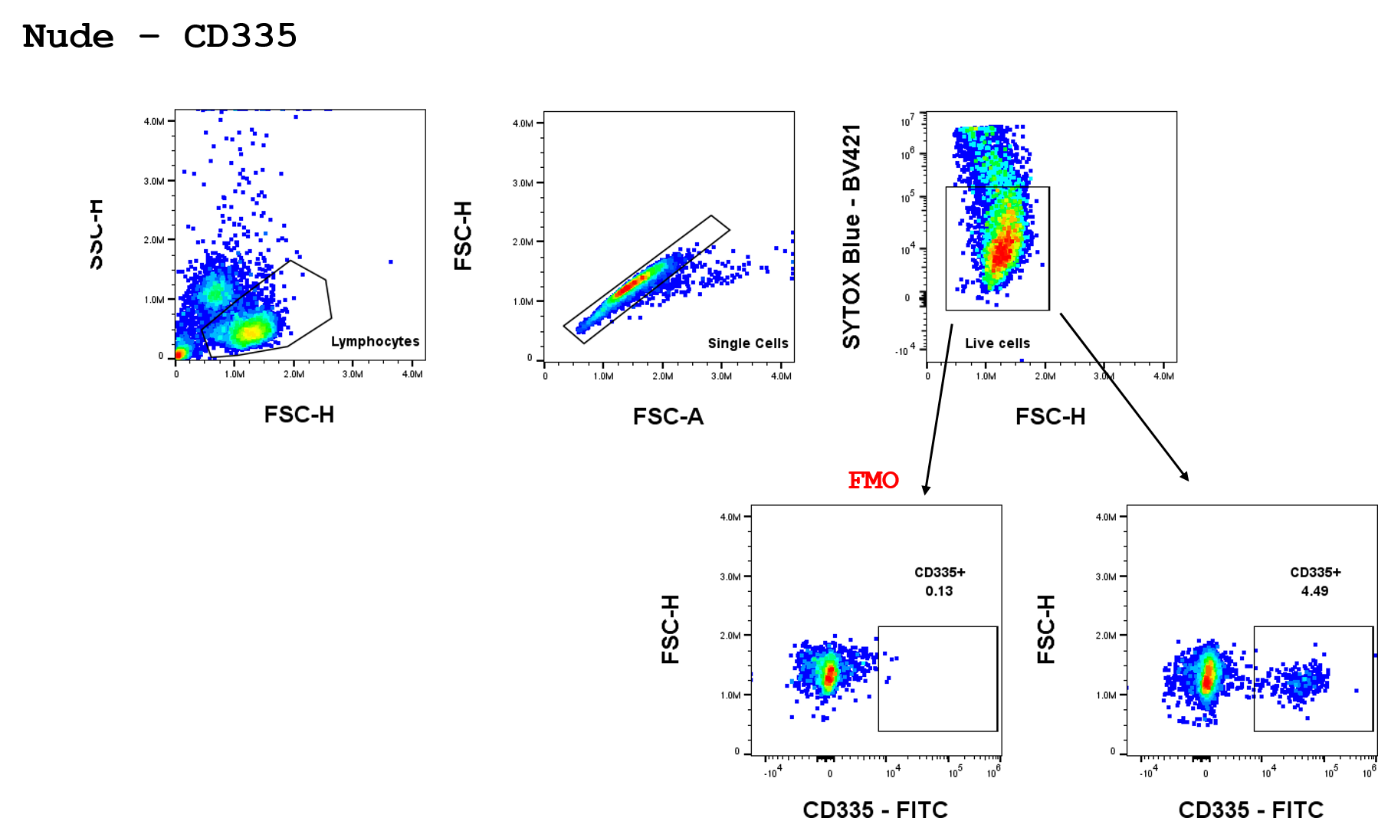

*Figure 3. Gating strategy for flow cytometric detection of CD335⁺ cells in PBMCs from nude rats.*

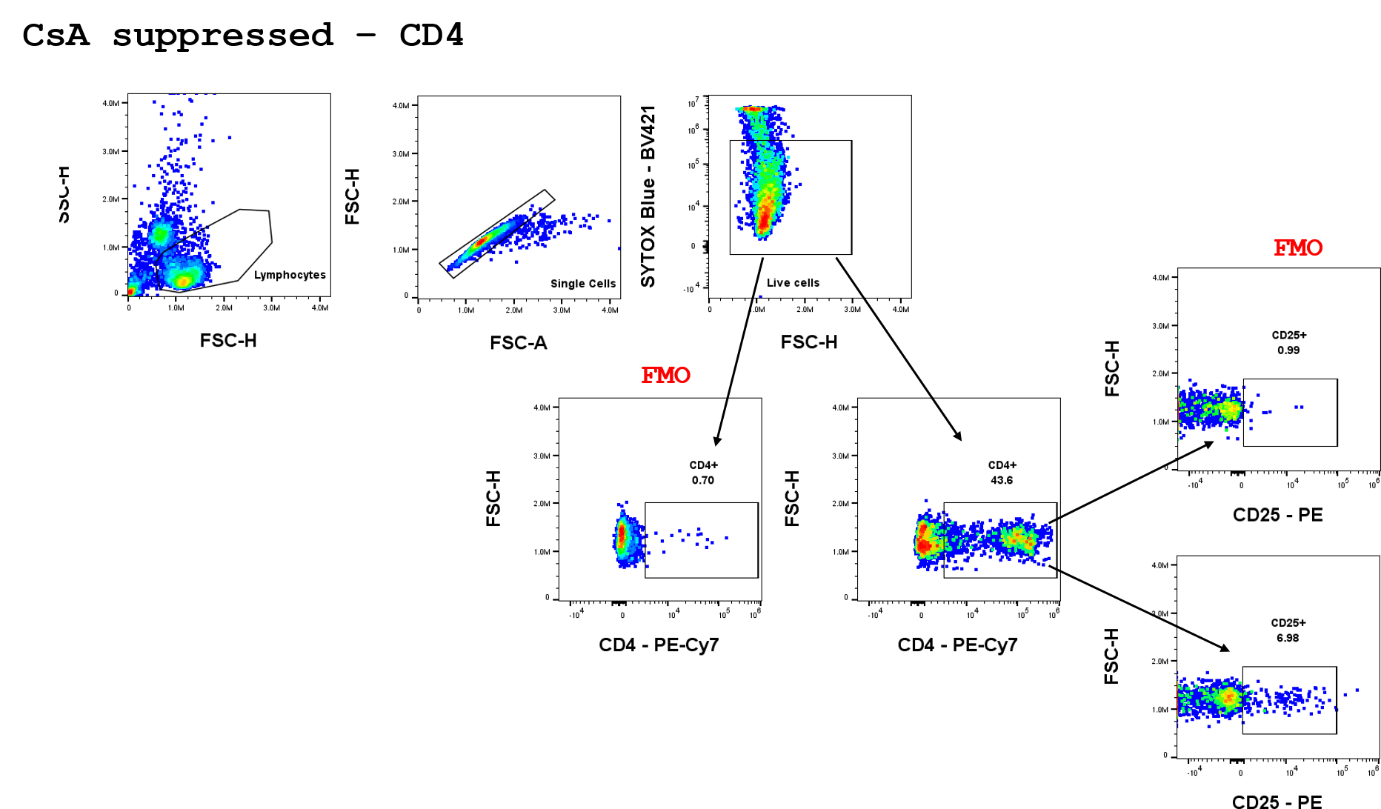

*Figure 4. Gating strategy for flow cytometric detection of CD4⁺ cells in PBMCs from CsA immunosuppressed rats.*

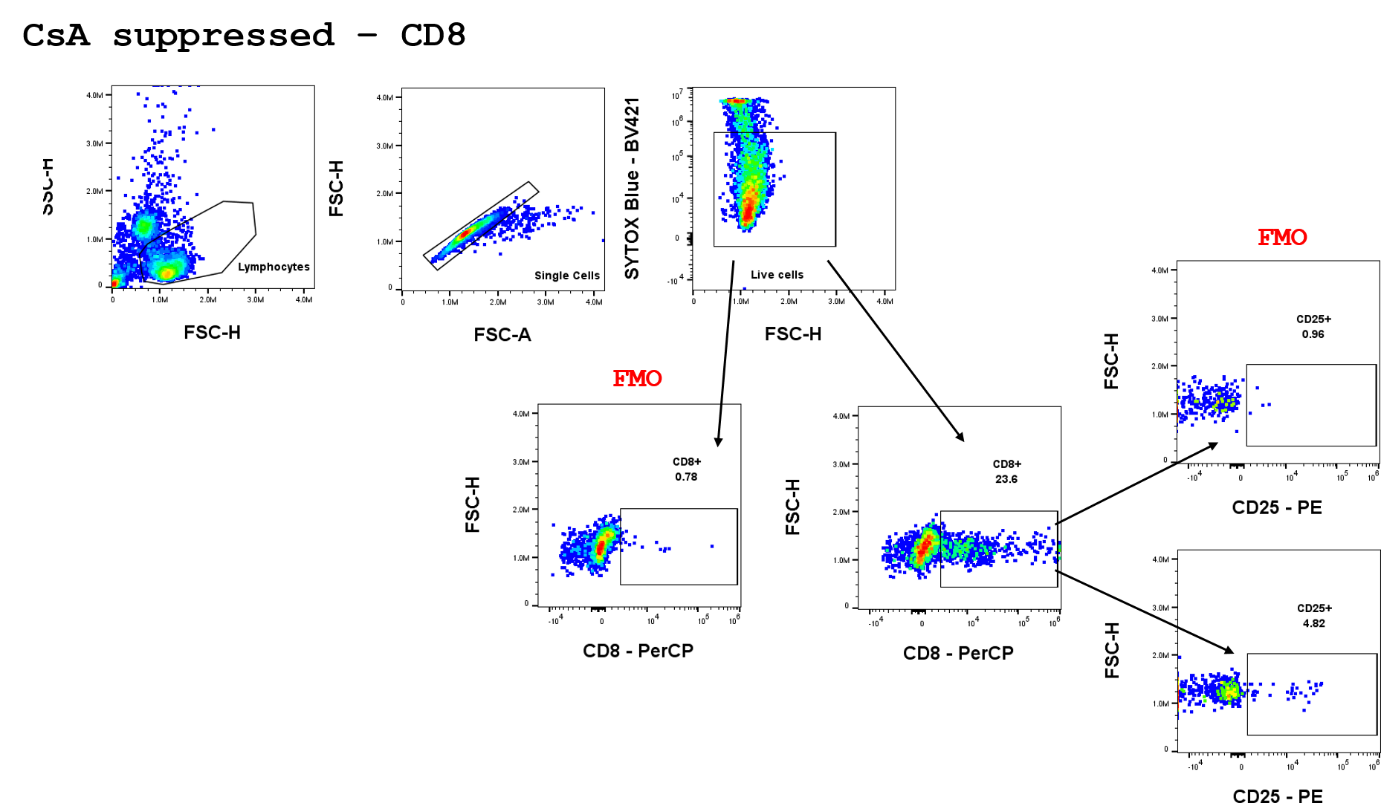

*Figure 5. Gating strategy for flow cytometric detection of CD8⁺ cells in PBMCs from CsA immunosuppressed rats.*

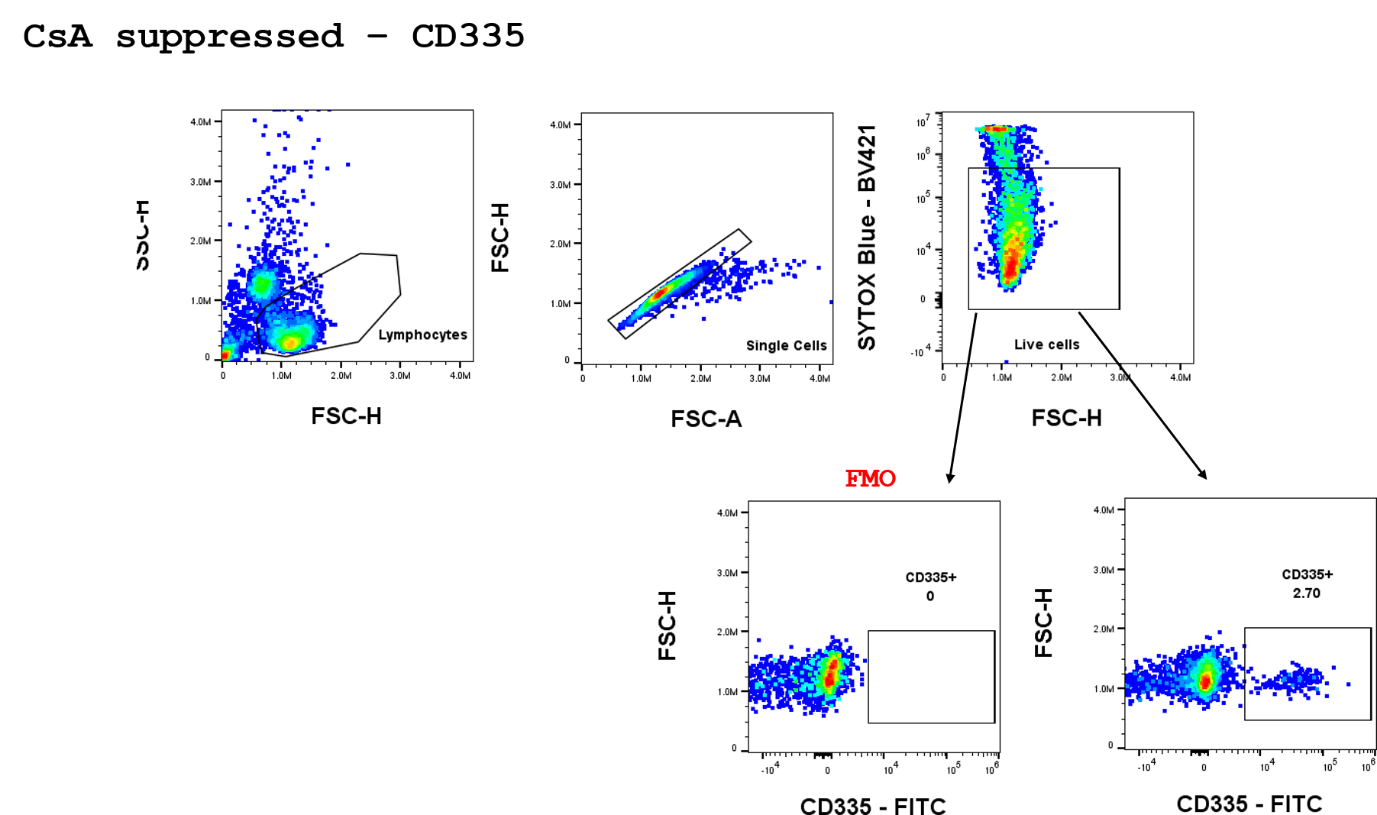

*Figure 6. Gating strategy for flow cytometric detection of CD335⁺ cells in PBMCs from CsA immunosuppressed rats.*
